# Heterosis across environmental and genetic space

**DOI:** 10.1101/2024.09.06.611759

**Authors:** Gabrielle D. Sandstedt, Catherine A. Rushworth

## Abstract

When genetically divergent lineages meet again in secondary contact, hybrids may suffer negative, fitness-reducing consequences, or benefit from positive genetic interactions that result in increased fitness. Empirical studies of heterosis, a phenomenon in which hybrids outperform their inbred progenitors, are of great interest in agriculture, but are less often performed in wild systems. In this study, we leverage *Boechera retrofracta*, a primarily self-fertilizing wildflower species, to explore how population divergence influences fitness effects upon secondary contact. We integrated genomic data and a large-scale fitness experiment to compare fitness and heterozygosity between outbred and inbred progeny of *B. retrofracta*. We show that interpopulation hybrids have increased overwintering survival compared to inbred individuals, indicative of heterosis. The magnitude of heterosis varied across genotypes and different environments, with overwintering survival increasing with genetic distance between parents. Sliding window analyses of genotyping by sequencing data show that heterozygosity varies across the genome of two species, *B. retrofracta* and the commonly co-occurring species *Boechera stricta*. We next compared these data with *de novo* F_2_s (intrapopulation, interpopulation, and interspecific crosses), as well as with wild-collected interpopulation cross *B. retrofracta* and interspecific *B. stricta* x *B. retrofracta* hybrids. Wild-collected interspecific hybrids appear to be F_1_s, while wild-collected intraspecific *B. retrofracta* are consistent with more complex crossing patterns. Because outcrossing is associated with a transition to asexuality in this group, this suggests different mechanisms underlie asexuality in hybrid and non-hybrid lineages. These findings underscore the potential differences in the role of heterosis between genetic groups at different stages of divergence and its relevance following hybridization in nature.

## Introduction

The evolutionary processes driving population divergence are dynamic over space and time. Thus, when populations come into secondary contact, the fitness of their hybrid offspring may deviate from that of progeny originating within populations. For instance, within populations, inbreeding depression can arise when mating of closely related individuals or self-fertilization reduces fitness due to increased homozygosity at deleterious recessive loci (Charlesworth & Charlesworth 1987; Ralls et al. 1988; Lynch 1989). Consequently, inter-population hybrids may exhibit increased fitness compared to individuals resulting from local inbreeding, a phenomenon known as hybrid vigor or heterosis. Though other explanations have been explored, heterosis seems most often to involve masking of deleterious recessive alleles through allele-allele interactions (i.e., dominance; Crow 1948, 1952; Lande and Schemske 1985, Charlesworth and Willis 2009). Heterosis has sparked substantial attention from researchers due to the insights it offers into the accumulation of deleterious mutations, or genetic load, within natural populations (Crow 1948; Lynch 1991; Whitlock et al. 2000), as well as its practical implications in agriculture (Darwin 1876; Shull 1952; Falconer 1981). Alternatively, outbreeding depression can emerge in hybrids between distant populations of the same species, reducing fitness and potentially balancing the effects of heterosis through genetic interactions (Edmands 1999; Dobzhansky 1951; Muller 1942; Lynch 1991).

The interplay between heterosis and outbreeding depression and its effects on hybrid fitness is shaped, in part, by the underlying genetic mechanisms. For example, the genetic load or accumulation of deleterious alleles within populations, which is masked in hybrids, may to a large extent determine the strength of heterosis. While theory suggests that the input of strongly deleterious mutations can be balanced by the rate of natural selection removing them (i.e., mutation – selection balance), their accumulation contributes to genetic load when selection is inefficient (Haldane 1937; Crow and Kimura 1970). Indeed, genetic load can be influenced by the degree of dominance of the deleterious mutations, mating system, and effective population size (Kimura et al. 1963; Lynch and Gabriel 1990; Lynch et al. 1995; Charlesworth, Charlesworth, & Morgan 1990). Even in populations that are predicted to effectively remove or “purge” lethal deleterious recessives, like populations that expose these mutations through self-fertilization (Lande & Schemske 1985), genetic load can persist with inbreeding when deleterious alleles have weak enough effects to evade selection (Charlesworth, Morgan, & Charlesworth 1990; Glémin 2003). Therefore, hybrid fitness should be enhanced in crosses that mask more of these deleterious recessive alleles, such as in diploid F_1_ hybrids that inherit a haploid genome from each parent, or with increasing genetic distance between parents (Lynch 1991; Edmands 1999; Edmands 2002). Still, hybrid fitness may instead be reduced by incompatible epistatic interactions that manifest either in the F_1_ or F_2_ generation (Bateson Dobzhansky Muller incompatibilities, BDMIs; Bateson 1909; Dobzhansky 1951; Muller 1942). In reality, hybrid fitness will reflect a combination of these dominant and epistatic interactions, resulting in a genomic mosaic underlying substantial variation in ecologically-relevant traits.

The environment is another factor known to influence fitness across the lifecycle. In particular, as populations diverge, they may adapt to different ecological niches (Sobel et al. 2010). Consequently, crosses between populations may lead to hybrid offspring that are maladapted to either of their parental habitats, resulting in a fitness cost (Coyne & Orr 2004). Recent studies have also highlighted the potential for negative epistatic interactions to shape fitness outcomes in response to environmental context (Thompson et al. 2023). These interactions are predicted to result from mismatched combinations of hybrid traits underlain by additive BDMIs (Thompson et al. 2023), though the presence of dominant BDMIs would predict different outcomes. On the other hand, hybrids may outperform parents either by displaying increased stability across environments or by capitalizing on favorable environments compared to inbred progeny (Cheptou & Donohue 2011). If hybrids show less variability in fitness across different environments – a phenomenon attributed to the inherent heterozygosity of a hybrid genome (i.e., “buffering theory”; Allard & Bradshaw 1964) – this enhanced stability could prove to be advantageous, especially under extreme conditions when hybrid performance is only mildly affected by stress. Other evidence has instead suggested that hybrids might show greater variability in fitness in different conditions, which may enable them to be more robust to changing environments (Mercer et al. 2007; Arnold & Martin 2010). Considering that stress can exacerbate inbreeding depression, the magnitude of heterosis is expected to vary across environments, potentially increasing in stressful conditions (reviewed in Cheptou & Donohue 2011). However, our current understanding of heterosis largely stems from studies in crops under extreme conditions (Li et al. 2004), leaving a gap in our understanding regarding the mechanisms of heterosis in nature and its broader implications for hybridization.

In this study, we leveraged the frequency of hybridization in the mustard genus *Boechera* Á. Löve & D. Löve (“rock cress”, formerly *Arabis* L.; Brassicaceae), an ecological model system native to North America (Rushworth et al. 2011, 2022). We focus on *Boechera retrofracta*, a widespread species that hybridizes with many congeners and primarily occurs in semi-arid sagebrush scrub and high desert (Windham and Al-Shehbaz 2006). Despite most of the 70+ sexual diploid *Boechera* species being highly self-fertilizing, intra- and interspecific hybridization is common (Song and Mitchell-Olds 2007; Schranz et al. 2005; Li et al. 2017). In the wild, hybrids are almost always asexual, consistent with a link between hybridization and a shift to apomixis, a process wherein asexual reproduction occurs via seed (Beck et al. 2012). Yet, controlled crosses almost never result in asexuality (for example, Schranz et al. 2005, Schranz et al. 2006, Rushworth and Mitchell-Olds 2021, this paper; but see Mau et al. 2021). The timing and molecular mechanisms of this transition are of key importance to understanding the origins and persistence of asexuality in this group. Previous studies suggest the transmission of apomixis through crosses between sexual and asexual lineages (Mau et al. 2021) and a strong association of alleles at two loci with apomixis (Mau et al. 2013; Corral et al. 2013). However, additional work is needed to confirm a causative relationship between these alleles and asexuality, and substantial effort will be necessary to characterize long-term dynamics of asexual alleles in natural environments.

Previous work has shown that wild interspecific and interpopulation crosses involving *B. retrofracta* have higher overwintering survival than sexual, self-fertilized *B. retrofracta*, but are more susceptible to insect herbivory later in the growing season (Rushworth et al. 2020). This is consistent with variable hybridization-induced fitness consequences across different traits, but the extent to which heterosis impacts over-winter survival remains unclear. Here, we investigate the consequences of population divergence on hybridization within *B. retrofracta*, combining a large-scale field experiment with genomic data from de novo F_2_s, wild-collected asexual B. retrofracta, and both de novo F_2_ and wild-collected asexual B. *stricta* x B. *retrofracta*. We leverage this approach to address the following questions: 1) Are survival and other fitness traits influenced by heterosis or outbreeding depression in crosses between divergent *B. retrofracta* populations? 2) Does the extent of parental divergence influence trait expression? 3) How do the environment and genetic composition of parents impact these traits? 4) Is heterozygosity variable across the genome of newly-formed interspecific and intraspecific crosses, and how does this pattern compare with wild-collected interspecific and intraspecific lineages? 5) How is the quantity of heterozygosity, and ultimately trait expression, of wild lineages affected by the timing of the transition to asexuality? Our findings revealed evidence of heterosis primarily in overwintering survival among interpopulation hybrids in *B. retrofracta*. We observed that the strength of heterosis in survival varies by genotype and environment, with increased parental divergence predicting greater survival.

Additionally, the transition to asexuality in hybrids can occur in later generations between populations of *B. retrofracta*, differing from the apparently fixed F_1_ transition in *B. stricta* x *B. retrofracta* hybrids. Finally, we discuss the role of heterosis, with regards to its influence on hybrid success, at different stages of species divergence.

## Methods

### PLANT GROWTH AND CROSSING

In fall 2012, lineages from existing, previously genotyped seed stocks (Song et al. 2006, Lee and Mitchell-Olds 2011, Rushworth et al. 2018) were selected for germination and crossing. Three cohorts of one line each from 10 populations of *B. retrofracta* were selected for interpopulation (“among”) crosses, and four lines each from three populations of *B. retrofracta* were crossed with other individuals from the same population to form intrapopulation (“within”) crosses (Table S1). Concurrently, ten lines of *B. stricta* were crossed with *B. retrofracta* (Rushworth and Mitchell-Olds 2021). All growth and germination was carried out as in Rushworth and Mitchell-Olds (2021). Briefly, seeds were germinated on wet filter paper, and seedlings were transplanted and grown in the Duke University Greenhouses until reaching maturity, at which point they were subjected to a 6-week vernalization period at 4C under 12-hour days to induce flowering. At the end of this period, plants were transferred to the Duke University Phytotron, where they were grown in two modified adjacent M13 growth chambers (Environmental Growth Chambers, Chagrin Falls, OH, USA) until flowering completed. Reciprocal crosses were conducted on 1-4 flowers among each potential pair of parents. Crosses were performed in late morning and early afternoon, just prior to peak anther dehiscence, which occurred approximately between 2 and 4pm (C. Rushworth, personal observation) following Schranz et al. (2005). Dry seed was collected upon maturation of all crosses that produced fruits. F_1_ seed was germinated, plants were grown to maturity in the Duke University Greenhouses, and tissue was collected for genotyping at three microsatellite markers (ICE3, c8, and BF20; Rushworth et al. 2018) to confirm crossing success (Table S2). Markers were selected based on known heterozygosity within *B. retrofracta*, ensuring that successful F_1_s would be heterozygous. All successful crosses were permitted to autonomously self-fertilize and set F_2_ seed. Fresh leaf tissue for GBS was collected on greenhouse-grown full siblings of F_2_s used in the field experiment, below.

### FIELD EXPERIMENT

In fall 2013, selfed seeds from parental lines and F_2_s were germinated as above, and plants were grown to maturity in the Duke University Greenhouses. Lines included 15 “among” F_2_s, 10 “within” F_2_s, and 21 parental lines (10 “among” and 11 “within”). 2000 plants were organized into randomized replicated blocks. At six weeks of age, rosette width was measured, and plants were shipped to central Idaho for planting into three experimental field sites. At one garden site (ALD), nearly all plants died following winter 2013/2014; thus, these data were removed from analyses (Fig. S1). Censusing was conducted upon multiple visits to each garden in the following spring and summer of 2014. On each plant we measured the following traits: life history stage, plant height, plant width, number of rosettes, number of bolting stalks, leaf number, fruit number, aborted fruit number, and two measures of insect herbivory: meristem removal and leaf damage as in Prasad et al. (2012). At the end of the season, one or two of the longest mature fruits were collected from 2 - 7 replicates of each genotype in each garden. We counted seeds in each fruit and calculated an averaged number of seeds for each genotype in each garden.

Averages were calculated for both viable-only and all (viable + inviable) seeds, as identified by eye. To obtain an estimate of fecundity (average maximum seed set per fruit), we multiplied this average by the number of fruits for each individual plant. Total fitness was calculated as survival * number of fruits * the average seed set per genotype per garden. *B. retrofracta* is known to be a short-lived perennial plant, but the vast majority of experimental plants do not survive or reproduce past their first year (Rushworth et al. 2020). Thus, fitness that was estimated in experimental year one closely approximates total lifetime fitness.

### STATISTICAL ANALYSES

We modeled the effects of line type, which we call “cross type” (*CT*; a factor that takes the value of within *B. retrofracta* populations or among *B. retrofracta* populations), garden (*garden*; a factor that takes on the value of CAP or MIL garden sites), and their interaction on fitness components across the lifecycle. Inbred parents were not statistically different from within population crosses across both gardens for any trait; thus, for most analyses, parents and within-population crosses were considered together (Table S3, S4). The model used is shown in the following equation, where *y* represents a fitness component, with initial plant width, block, position within block, and genetic relatedness (see below) incorporated as random effects:

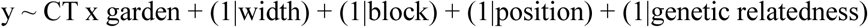

To model these effects on fitness, we performed Bayesian Regression Models using Stan (brms) in the R Statistical Software (v4.2.2; R Core Team 2022; Bürkner 2017), which provides a flexible framework for estimating Bayesian models and addresses complexities of model specification. In brms, we used the brm() function with default priors and an appropriate family distribution for the fitness trait analyzed. We fit survival, reproduction, flowering, and meristem removal with a bernoulli distribution; relative fruit number, fruit number * average seed set, stalk number, plant height, plant width, longest stem, and leaf number with a skew_normal distribution; leaf herbivory (“damage”) with hurdle_gamma distribution; aborted fruit number with a hurdle_poisson distribution; and rosette number and total fitness with a hurdle_lognormal distribution. Models for survival and total fitness included all plants scored in the field (*N* = 1,229), whereas models for all other traits only included plants that survived (*N =* 780).

We also utilized brms to estimate family-level mean survival for each garden separately. These models included family as a fixed effect, block as a random effect, and initial plant width as a covariate (as shown below) and were fit with a bernoulli distribution:

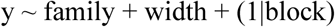

We ran each model with two to four chains of 2000-4000 iterations per chain, and a warm-up of 1000 iterations. We rigorously assessed model fit using several approaches in brms: we compared the distribution of simulated data from posterior predictive checks to observed data using the pp_check() function, we examined convergence with summary() and ensured that bulk and tail effective sample sizes > 1000 and the potential scale reduction factor (R^) ∼ 1.00, and we performed model comparison using the leave-one-out (loo) function. For each model output, we visualized conditional effects with 95% confidence intervals of cross type and garden on each trait using the conditional_effects() function.

Finally, we used the hypothesis() function to conduct post-hoc comparisons between effects of cross types at each garden site, which estimates differences when the posterior probability exceeds 95% confidence.

### GENERATING AND PROCESSING GENETIC DATA

We prepared 190 samples for Genotype by Sequencing (GBS), including 22 sexual *B. retrofracta* individuals (N = 1 – 5 individuals from 14 populations = 22), 31 F_2_s derived from intra-population crosses of *B. retrofracta* (N = 3 – 4 individuals per 10 unique cross combinations as above = 31), 46 F_2_s derived from inter-population crosses of *B. retrofracta* (N = 3 – 4 individuals per 15 unique cross combinations as above = 46), and 28 wild-collected asexual *B. retrofracta* x *B. retrofracta* hybrids (N = 2 replicates of 1 individual per 14 populations = 28; Table S2). For comparison with another naturally hybridizing pair involving *B. retrofracta*, we incorporated five *B. stricta* individuals (N = 1 individual each per 5 populations = 5), 22 F_2_s derived from crosses between species of *B. stricta* and *B. retrofracta* (N = 3 – 7 individuals per 4 unique cross combinations = 22; Rushworth and Mitchell-Olds 2020), and 32 wild-collected asexual *B. stricta* x *B. retrofracta* hybrids (N = 2 replicates of 1 individual per 16 populations = 32; Table S2). We determined that four natural hybrid asexuals (N = 2 replicates of 1 individual from 2 populations = 4) did not belong to our focal species and thus later removed them from downstream analyses, resulting in 186 samples. We extracted DNA from leaf tissue from all plants using the Qiagen DNeasy® Plant Mini Kit and protocol (Qiagen, Carlsbad CA). We submitted samples to the Cornell Genomics Facility, where genomic DNA was digested using the PstI restriction enzyme. The resulting fragments were sequenced on the Illumina HiSeq 2500 platform to generate 96-bp single-end reads.

To process GBS data for 186 samples, we prepared raw sequences for alignment by removing Illumina and unique adapters and discarding reads shorter than 50-bp with Cutadapt (Martin 2011; Table S2). We confirmed removal of adapters using FastQC (Andrews 2010). Given that two replicates represented the same hybrid asexual sample, which should be identical, we concatenated these FASTQ files prior to alignment, resulting in 156 samples. We aligned single-end reads to the *B. stricta*, LTM version_2 reference genome using BWA *mem* (Li and Durbin 2009; Li 2013) and removed reads with an alignment quality below Q29 with SAMtools (Li et al. 2009). To prepare files for variant calling, we added read groups with AddOrReplaceReadGroups using Picard tools (http://broadinstitute.github.io/picard). To generate variant and invariant sites for all samples, we used Genome Analysis Toolkit’s (GATK) HaplotypeCaller, combined all samples using GenomicsDBImport, and performed joint genotyping with the the “-all-sites” option using GenotypeGVCFs (McKenna et al. 2010). We split this initial Variant Call Format (VCF) into two using GATK’s SelectVariants: one contained biallelic single nucleotide polymorphisms (SNPs) and another contained invariant sites only. To filter for high-quality genotypes, we filtered both variant and invariant VCFs using GATK’s VariantFiltration, which removed sites with a quality score (QUAL) < 30, root mean square mapping quality (MQ) < 40, mapping quality rank sum test (MQRankSum) < −12.5, read position rank sum test (ReadPosRankSum) < −8, and depth (DP) < 5 and > 200. We further filtered the biallelic SNP VCF, excluding sites with a quality by depth (QD) score < 2, strand odds ratio (SOR) > 3, and fisher strand (FS) > 60. Additionally, we omitted heterozygous positions with a ratio of reference to alternative alleles < 25% or > 75% using the package “SNPfiltR” implemented in R (DeRaad 2023). Finally, we merged invariant (415,163 positions) and variant (27,679 positions) VCFs, referred to as the “merged VCF” below. However, most of these variant positions included rare alleles and low coverage across samples. Therefore, we excluded positions with a minor allele frequency (MAF) < 5% and removed sites that had called genotypes for <10% of the samples using VCFtools for each analysis – SNPs remaining after these filtering steps are reported in detail below (Danecek et al. 2011). We note that the count of invariant sites remained unchanged after filtering for <10% coverage at a position.

### Genetic Matrix

We incorporated a genetic relatedness matrix into our brms models to account for any genetic structure effects. To construct this matrix, we downsampled our merged VCF to include only crosses and parents from the field experiment. Given that there were three replicates per cross, we stochastically downsampled to a single replicate and excluded invariant sites with GATK’s SelectVariants (*N* = 42; SNPs = 2,064). Subsequently, we generated the final genetic relatedness matrix using PLINK (Purcell et al. 2007).

### Nucleotide diversity (d_XY_) between *B. retrofracta* parents

To explore the relationship between fitness and genetic divergence of *B. retrofracta* parents involved in crosses, we downsampled the merged VCF to only include sexual parental genotypes (*N* = 21; SNPs = 2,263; Table S1). We used *Pixy* to calculate divergence (d_XY_; Korunes and Samuk, 2021) across non-overlapping 1Mbp windows, and we obtained genome-wide values by computing the ratio of count differences to count comparisons. We modeled the relationship between d_XY_ and family-level means in survival with a generalized linear model (GLM) using the glm function in the R package “lme4” (Survival ∼ d_XY_; Bates et al. 2007) and conducted an ANOVA utilizing the anova function in the “car” package with type III sums of squares.

### Assessing patterns of genomic variation

To visualize patterns of genetic relatedness across the 156 *Boechera* samples corresponding to the following six groups: sexual *B. retrofracta*, sexual *B. stricta*, *de novo* sexual F_2_ hybrids and asexual wild-collected hybrids (*B. retrofracta* x *B. retrofracta*, *B. strica* x *B. retrofracta*), we utilized our merged VCF containing 3,862 biallelic SNPs after filtering. We calculated sequence diversity within groups (π) and pairwise divergence between groups (d_XY_; *Pixy*: Korunes and Samuk, 2021). In addition, we used a custom python script to scan the merged VCF, calculating the number of heterozygous positions (*i.e.,* 0/1 in a VCF) in 5Mb windows for each sample and averaged heterozygosity across the six groups to determine whether heterozygosity was localized to particular regions of the genome. Furthermore, we assessed patterns of genomic variation using a principal components analysis (PCA). For this analysis, we exclusively considered SNP positions, excluding invariant sites, and conducted the PCA with the R package “SNPRelate”.

Moreover, we calculated the average genome-wide admixture proportion and global ancestry for individuals using the model, ENTROPY (Shastry et al. 2021). This model implements a hierarchical Bayesian approach that leverages genotype likelihoods to inform genotype and ancestry parameters. Our analyses focused on two hybridizing pairs: 1) populations of *B. retrofracta*, and 2) *B. stricta* and *B. retrofracta*. For each hybridizing pair, we generated genotype-likelihood files from downsampled VCFs to include only the respective intra-specific species, *de novo* sexual F_2_s, and asexual hybrids (*B. retrofracta* populations and hybrids = 113 samples, 2,134 SNPs; *B. stricta* and *B. retrofracta* populations and hybrids = 65 samples, 3,819 SNPs). We ran ENTROPY with the same model parameters across three chains for both hybridizing pairs. These parameters included setting putative demes (-k) to two, inter-specific ancestry (-Q) to one, MCMC steps (-l) to 15,000, burn-in (-b) to 5000, storing every step after burn-in (-t) at five, and assigning a scalar for Dirichlet initialization of q (-s) to 30. We assessed convergence of these models, ensuring that R^ ≈ 1 and effective sample sizes > 1000 for each individual. Finally, we extracted admixture proportion (q) and ancestry (Q) using the “estpost.entropy” function with the “-p” option and visualized these values with ggplot2 in R (Wickham 2011).

## Results

### ELEVATED SURVIVAL IN OUTCROSSED LINEAGES

To determine the fitness consequences of putative divergence among *B. retrofracta* populations, we compared 15 fitness traits across the lifecycle in individuals produced by local inbreeding (self-fertilized or within population crosses) with those from outcrossed lineages (among population crosses) in two common gardens, CAP and MIL. One crucial fitness trait stood out: outcrossed lineages had higher overwintering survival than offspring produced by local inbreeding at both CAP (within = 0.260.16, among = 0.480.20) and MIL (within = 0.750.15, among = 0.880.08; Fig 2a; Table S5, S6), indicative of heterosis. The only other trait that showed potential evidence for heterosis was plant height; however, elevated plant height in outcrossed lineages was driven only by the effect at the CAP garden (within = 165.9017.27, among =183.5117.37) and not MIL (within = 103.4915.82, among = 117.40; Fig S2; Table S5, S6, S7). Despite increased survival in outcrossed lineages for both gardens, total fitness was not impacted by cross type (Fig. 2b; Table S5, S6). Instead, we only observed a garden effect on total fitness, with plants at the CAP garden exhibiting increased total fitness (within = 269.5729.15, among = 323.1734.86) compared to plants at the MIL garden (within =146.9016.05, among = 148.6214.19; Fig 2b; Table S5, S7). Herbivory did not differ between cross type in either garden (CAP: within = 0.0250.003, among = 0.0260.003; MIL: within = 0.0270.003, among = 0.0310.003; Fig. S2, S3; Table S5, S6). These findings suggest that beyond survival, heterosis was generally modest across both gardens. In addition, fitness was not decreased in among-population crosses for any observed trait, consistent with a lack of outbreeding depression in our data (Fig. S2, S3; Table S5, S6).

**Figure 1:**
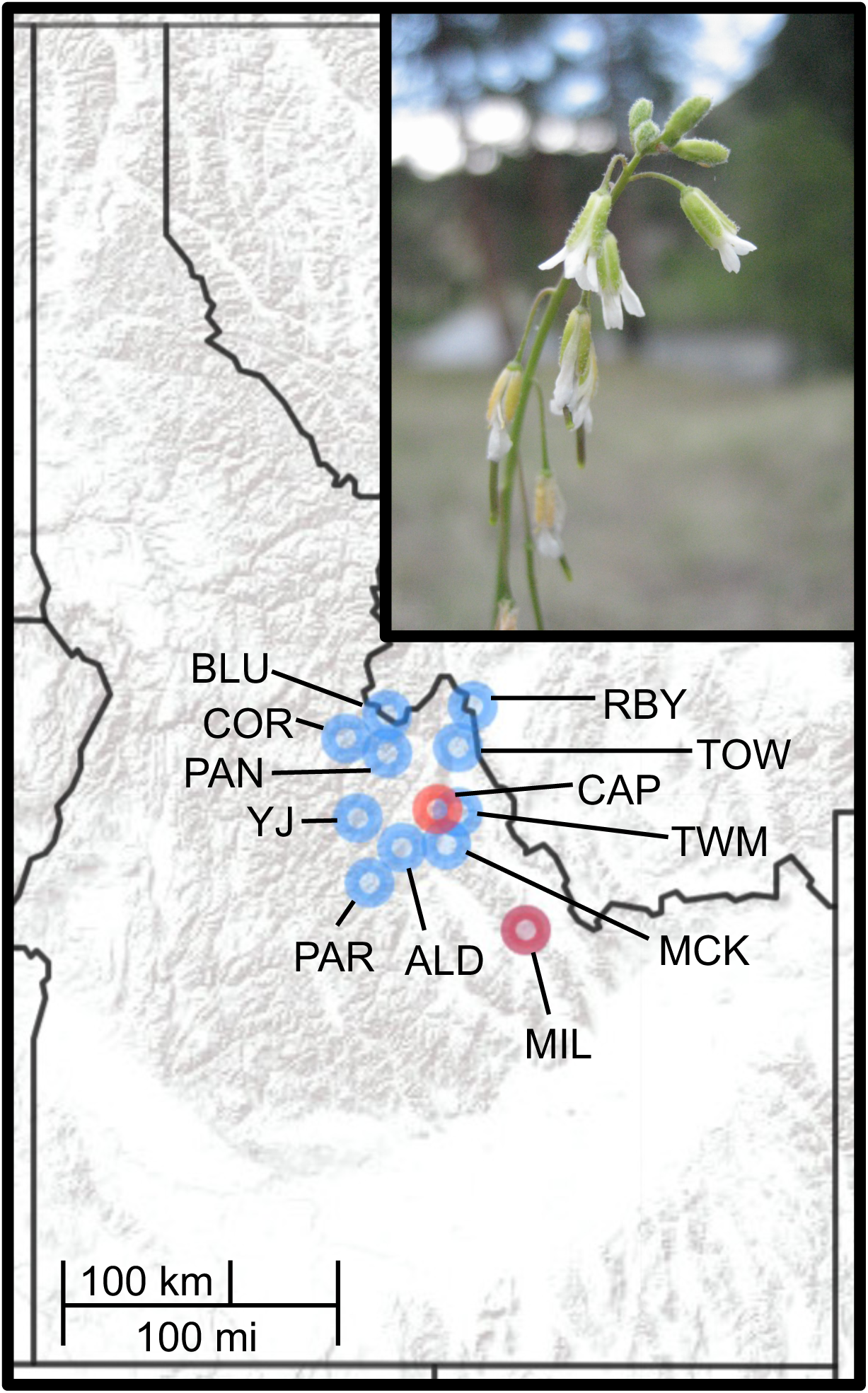
Distribution map of Idaho indicating *B. retrofracta* populations used in this study and the two common garden sites. Population identity is shown as blue points with a population code. Two common garden sites designated by red points (CAP and MIL). Nearly all plants died at a third garden site (ALD) and was removed from analyses. Photograph of a *B. retrofracta* plant in top right panel, captured by Dr. Rushworth.

**Figure 2:**
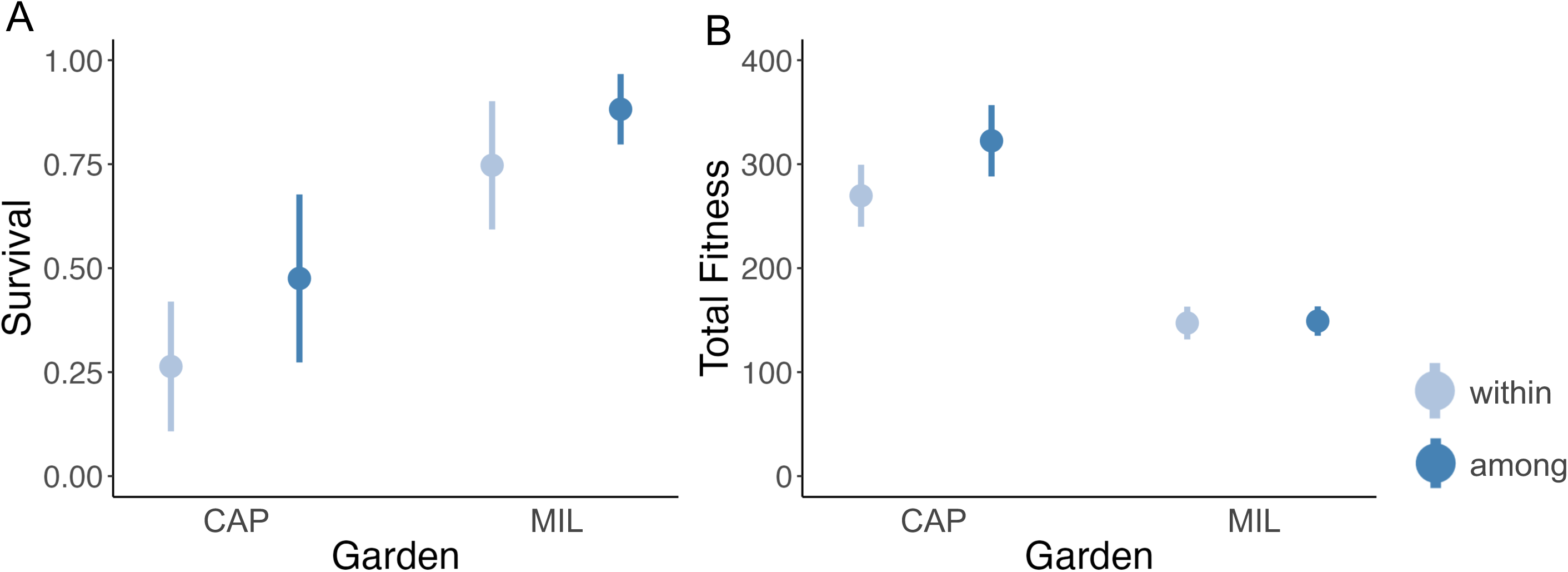
Heterosis is primarily expressed as survival across two common gardens, CAP and MIL, in *B. retrofracta*. The data points depict conditional means of fitness traits of individuals within (light blue) and among (dark blue) populations, with standard error bars shown. **A)** Survival is increased in among populations. This trait was assessed based on whether individuals did not survive (0) or survived (1). **B)** Total Fitness shows no evidence for heterosis or outbreeding depression. Total fitness was calculated as the product of survival, number of fruits and average seed set per genotype per garden.

We next investigated the relationship between parental genetic divergence and family-level mean survival. We hypothesized that increased genetic distance between parental pairs results from a larger quantity of divergent alleles, which are predicted to be recessive and weakly to strongly deleterious (Ohta 1973). The process of divergence thus provides increased opportunities for masking of such recessive alleles, leading to higher rates of survival. Our results show a positive correlation between greater genetic distance between parents and increased survival in both gardens (CAP: χ2 = 9.23, df = 1, *P* = 2.38 × 10^−3^; MIL: χ2 = 11.50, df = 1, *P* = 6.97 × 10^−4^; Fig. 3; Table S1, S8). We also note here that divergence does not appear to be a smooth function, with two apparent groups in the among-population crosses. One group clustered near a d_XY_ of 0.0010 and another near 0.0020 (dark blue data points, Fig. 3; Table S1), suggesting potential population structure within *B. retrofracta*.

**Figure 3:**
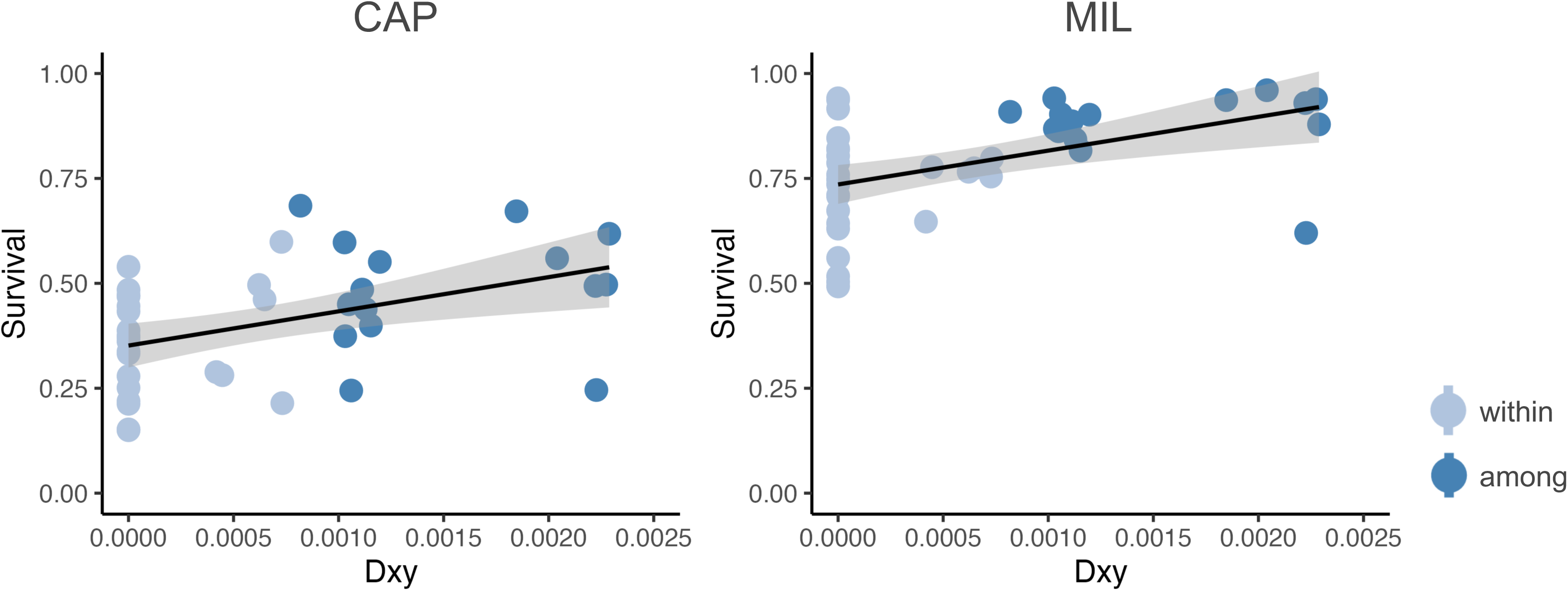
Survival increases with increasing genetic distance in both gardens for *B. retrofracta*. Left panel shows survival in CAP garden and right panel depicts survival in MIL garden. Data points represent conditional means for survival within (light blue) and among (dark blue) population cross combinations. X-axis represents divergence between parents of each cross combination – points falling on the 0 of the x-axis represent parents that self-fertilized. Data are fitted with a generalized linear model: survival ∼ Dxy. Gray shading represents the standard error in the model.

### THE STRENGTH OF HETEROSIS VARIES BY GENOTYPE AND ENVIRONMENT

We next used two approaches to measure the extent to which genotype and environment influenced the expression of heterosis in survival. First, **to determine whether the strength of heterosis varied across gardens**, we compared the difference in mean survival between among- and within-population crosses, relative to among-population mean survival at CAP and MIL (*W*_among_ *– W*_within_ / *W*_among_; Oakley and Winn 2012). Second, **to determine whether the strength of heterosis varied across different unique crossing combinations between gardens**, we compared the difference between family-level mean survival of six within-population cross combinations and 15 among-population cross combinations to their corresponding mid-parent averages, relative to mid-parent average survival for each garden (W_family-level mean_ – W_mid-parent average_/W_mid-parent average_; Lynch & Walsh 1998).

Our first approach revealed a substantial difference in the strength of heterosis in survival between our two gardens – heterosis was threefold greater in CAP than MIL (46% vs. 15%, respectively). While this difference suggests variability in the expression of heterosis between the gardens, it is important to note that our brms model did not reveal a significant effect of garden site on survival (Table S5, S7). In our second approach, we quantified relative survival for all cross combinations. For the six within population crosses, all but one (wi9 at CAP) overlapped with their mid-parent average (Fig. 4, Table S9). For the 15 among population crosses, seven (47%) surpassed their mid-parent average in relative survival in each garden. However, only four (27%) consistently demonstrated an increase in relative survival at both CAP and MIL, highlighting the environmentally-dependent nature of heterosis (Fig. 4, Table S9). Together, these findings underscore the importance of both genotype and environmental factors in shaping the expression of heterosis in *B. retrofracta*.

**Figure 4:**
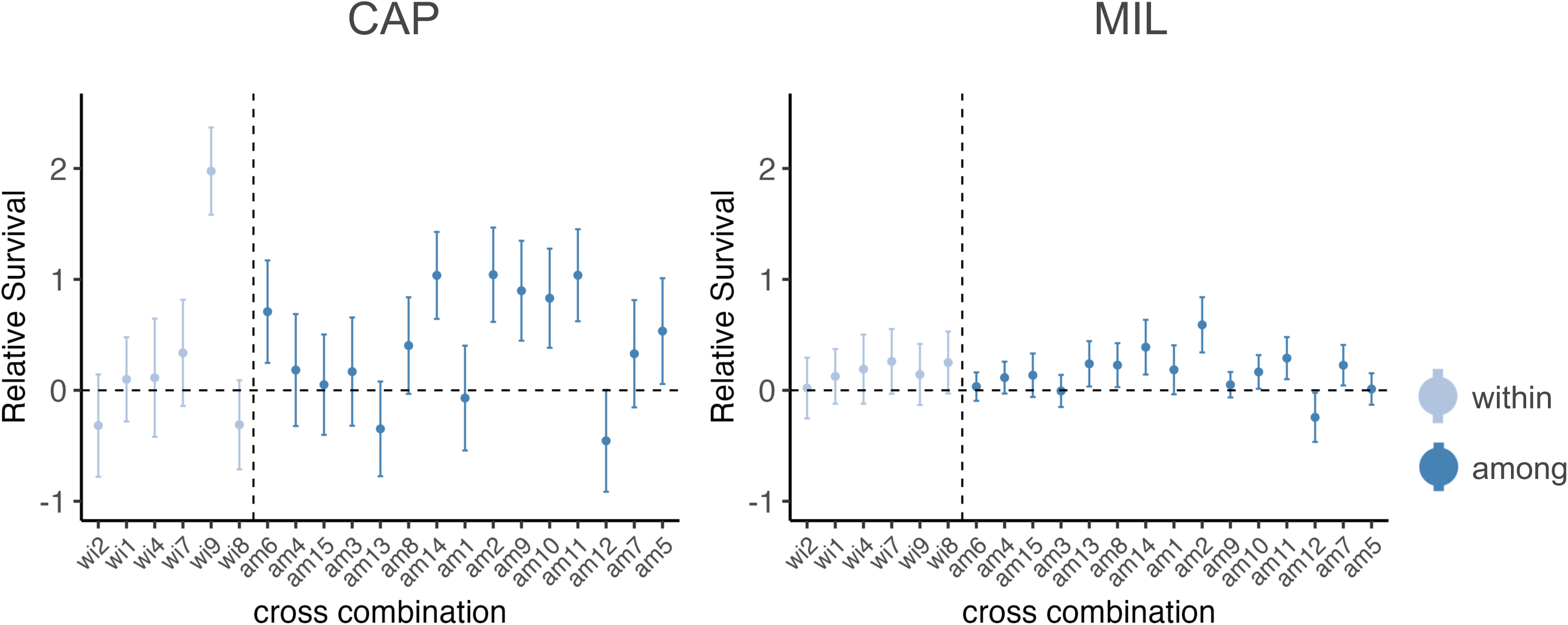
Heterosis is specific to genotype and environment in *B. retrofracta*. The left panel shows relative survival in CAP garden, while the right panel shows relative survival in MIL garden. Data points represent relative survival for six within (wi) and 15 among (am) cross combinations. Relative survival was calculated as the difference between conditional means for within and among cross combinations and their corresponding mid-parent averages, then standardized to their mid-parent average. Standard error bars were estimated using error propagation. Dashed horizonal line at 0 represents the standardized mid-parent average. Dashed vertical line separates within-population (light blue) and among-population (dark blue) cross combinations.

### GENETIC VARIATION AND HYBRIDIZATION IN WILD *BOECHERA*

Next, we explored patterns of genomic variation in parents, the two cross types, and wild-collected asexual *B. retrofracta* lineages. We aimed to compare the genomic diversity of these groups with interspecific hybrids, as both are frequently found in the native range of *Boechera*. Thus, we included both wild-collected asexual *B. stricta* x *B. retrofracta*, the most common hybrid parental combination in our study area (Rushworth et al. 2018), and *de novo* F_2_s between *B. stricta* and *B. retrofracta* (Rushworth & Mitchell-Olds 2021). Principal components analysis revealed substantial genetic structure among these comparison groups (Fig. 5A). The first principal component (PC1, 32.17%) explained interspecific variation between *B. stricta* and *B. retrofracta*, and the second (PC2, 13.71%) delineated genetic structure within *B. retrofracta*, with two distinct clusters of *B. retrofracta* separated by PC2 (Fig. 5A). As expected, both *de novo* and natural hybrids fell within and between their corresponding parental genetic groups (Fig. 5).

**Figure 5:**
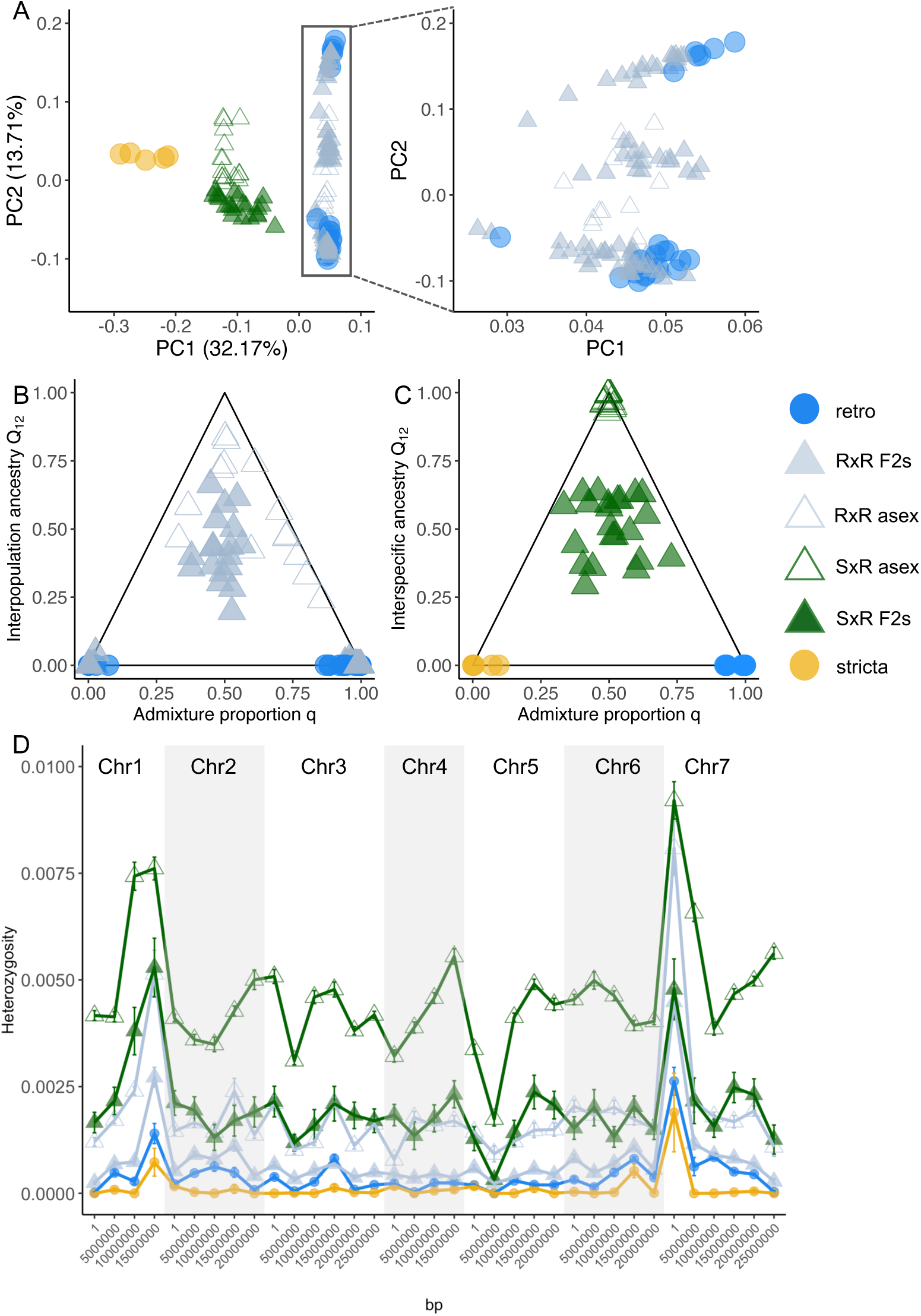
Genetic variation and hybridization in *Boechera*. Panels show *B. retrofracta* sexual parents (retro, blue circles), *de novo* sexual F2s derived from among and within *B. retrofracta* population crosses (RxR F2s, gray triangles), *B. retrofracta* asexual hybrids (RxR asex, gray outlined triangles), *B. stricta* sexual parents (stricta, yellow circles), *de novo* sexual F2 *B. stricta* x *B. retrofracta* hybrids (SxR F2s, green triangles), and *B. stricta* x *B. retrofracta* asexual hybrids (SxR asex, outlined green triangles). **A)** PCA of genetic relatedness among *Boechera* groups. The x-axis (PC1) accounts for 32.17% of the genetic variation and the y-axis explains 13.71% of the variation. Top right panel zooms into the genetic groups in *B. retrofracta*. **B)** Interpopulation ancestry (Q_12_) and admixture proportion (q) in *B. retrofracta*. **C)** Interspecific ancestry and admixture proportion between *B. stricta* and *B. retrofracta*. Solid lines represent the maximum values of ancestry. **D)** Heterozygosity in 5Mb windows along the seven chromosomes, with heterozygosity represented by the proportion of heterozygous positions (het / [het + hom]).

Strikingly, the two genetic clusters within *B. retrofracta* corresponded to the two distinct groups of among-population crosses identified using d_XY_ (Fig. 3, 5A). These qualitative observations were further supported by our quantitative analysis of genetic variation. To calculate divergence between these two *B. retrofracta* clusters, we assigned them to be distinct genetic groups in *Pixy*. The resulting π (0.006 and 0.008) and d_XY_ (0.0019) were modest (Fig 4). Crosses within genetic clusters had even lower divergence (d_XY_ = approximately 0.001; Fig 4), consistent with low divergence in the PCA (Fig. 5A). In contrast, the estimated divergence between *B. stricta* and *B. retrofracta* (d_XY_ = 0.0045) was over two-fold greater than between genetic subgroups of *B. retrofracta* (d_XY_ = 0.0019).

Despite the prevalence of intra- and interspecific asexual lineages in *Boechera*, little is known about the timing of the transition to asexuality. However, the stage at which asexuality takes effect profoundly impacts predictions of hybrid fitness and asexual longevity. Thus, we next estimated genome-wide admixture and global ancestry of our intraspecific and interspecific wild-collected asexuals, in comparison with all *de novo* crosses. Based on admixture proportion (q in Fig. 5), our ENTROPY model split *B. retrofracta* into two genetic groups (Fig. 5A). This mirrored the same genetic groups denoted in the PCA (Fig 3, Table S10) and supported by estimates of divergence. As expected, *de novo* F_2_s derived from crosses from between genetic groups vary around an admixture proportion and ancestry of 0.5 (Fig. 5B), where non-zero values of interpopulation or interspecific ancestry denote hybridization events (Q_12_ in Fig. 5B and 5C). F_2_s with an ancestry of 0 or admixture proportion of either 0 or 1 were also found, representing within population crosses or among population crosses from the same *B. retrofracta* genetic group (Fig. 5A). In natural asexual *B. retrofracta* interpopulation hybrids, there was considerable variation in admixture and ancestry – most appear to be later generation hybrids with greater shared ancestry with one *B. retrofracta* genetic group than the other (Fig. 5A and 5B, Table S10). This contrasts with natural asexual *B. stricta* x *B. retrofracta* hybrids, which displayed equal proportions of admixture and ancestry between the two parental species, consistent with the transition to asexuality occurring in the F_1_ generation (Fig. 5C, Table S11). We note that *de novo* F_2_s between *B. stricta* and *B. retrofracta* varied around an admixture proportion and ancestry of 0.5, similar to *B. retrofracta* F_2_s. These comparisons highlight how hybridization and the transition to asexuality manifest differently across *Boechera* species.

### GENOMIC LANDSCAPE OF HETEROZYGOSITY IN PARENTS AND HYBRIDS

Finally, we asked if any genomic patterns of heterozygosity might be identifiable in wild-collected interspecific and intraspecific hybrids. Since the single trait of survival exhibited heterosis, we might expect heterozygosity to be elevated in key regions of the genome underlying survival variation. Alternatively, survival is a highly polygenic trait (Paina et al. 2016), and thus heterozygosity may be elevated across many regions of the genome. We predicted that wild-collected hybrids, which have survived at least one cycle of natural selection in nature, might thus have a distinct pattern of heterozygosity reflecting their previously documented fitness advantage (Rushworth et al. 2020). Our sliding window approach revealed two elevated regions of heterozygosity: one along one arm of chromosome 1 and the other near the beginning of chromosome 7 (Fig 5D). Heterozygosity in these regions was elevated for all hybrids, whether interspecific or intraspecific, and whether lab-generated or wild-collected, and in both *B. retrofracta* and *B. stricta* selfed parental lines (Fig. 5D). *B. stricta* x *B. retrofracta* hybrids exhibited two regions of altered heterozygosity unique to this species combination but shared by *de novo* and wild-collected samples: a region of less sharply elevated heterozygosity at the end of chromosome 4, and a region of decreased heterozygosity located at the start of chromosome 5 (Fig. 5D). Surprisingly, the end of chromosome 1, start of chromosome 5, and start of chromosome 7 have been previously identified as regions of high genetic diversity in *B. stricta* (Wang et al. 2019).

## Discussion

In our study, we explored the consequences of population divergence on hybridization in a wild mustard species, *B*.*retrofracta*. We observed elevated overwinter survival in outcrossed lineages of *B. retrofracta* and found that its strength increased with genetic distance between parents, suggesting that hybridization masks deleterious alleles underlying survival. These findings were consistent with heterosis, with its expression varying by genotype and environment. Furthermore, we identified two distinct genetic subgroups within *B. retrofracta*, which represent the parents of most wild-collected *B. retrofracta* hybrids. These hybrids often transition to asexuality in later generations. This contrasted with wild-collected hybrids between the divergent species *B. stricta* and *B. retrofracta,* in which all asexuals appeared to be F_1_ hybrids. Below, we discuss these results in the context of the dual influence of genetic divergence and environmental variation on fitness to shape hybridization dynamics.

### The genetic underpinnings of heterosis

Heterosis has been a central topic in agricultural science due to its ability to counteract inbreeding depression in domesticated species (Shull 1952). The discovery in the 1930s that crossing elite maize lines produced F1 hybrids with improved yield and vigor marked a breakthrough (Duvick 2001). These hybrids often surpass their midparent values in traits such as yield, plant size, and survival (Springer & Stupar 2007). The exploration of heterosis has since expanded to other crops (Xiao et al. 1995; Krieger et al. 2010), model plant systems like *Arabidopsis* (Barth et al. 2003; Meyers et al. 2010; Dapp et al. 2015) and, more recently, non-model systems (Soto et al. 2023; Wang et al. 2024). Despite decades of research, the precise genetic basis of heterosis remains unclear. While most studies support the hypothesis that heterosis results from the masking of deleterious recessive alleles (Charlesworth & Willis 2009), others highlight the complexity involved (Liberatore et al. 2013; Huang et al. 2016) which may explain why a simple correlation between heterozygosity and fitness is not consistently observed, especially in wild species (Willi & Van Buskirk 2005; Oakley et al. 2019). Our study likewise does not reveal the precise genetic basis of heterosis. However, the combination of field and genomic data for *de novo* crosses and their selfed parents, along with genomic data from natural hybrids, offers powerful insight into the genetic mechanisms, as well as the ecological context, of heterosis in wild plants.

In our study, heterosis was observed in a single trait: over-winter survival (Fig. 2). This was perhaps surprising given the expectation that heterosis mainly arises from masking a large number of deleterious recessive alleles (Charlesworth and Willis 2009), suggesting they would be scattered across the genome. However, uneven expression of heterosis in specific traits has been frequently documented (Schemske 1983; Schoen 1983; Hamilton & Mitchell-Olds 1994; Husband & Schemske 1996), including in wild-collected *Boechera* hybrids (Rushworth et al. 2020). This might suggest that genetic load can concentrate in regions linked to variable traits, consistent with our finding that heterozygosity was elevated in two genomic regions for both intraspecific and interspecific F_2_ hybrids (chromosome 1 & 7 in Fig. 5).

Additionally, we observed that heterosis in overwinter survival was present across both experimental garden sites in F_2_ *Boechera retrofracta* crosses, which may be unexpected given that studies often report reduced fitness in the F_2_ generation (Sweigart et al. 2006; Stelkens et al. 2015; Zuellig & Sweigart 2018). F_1_ hybrids should exhibit maximized heterozygosity (and thus heterosis), but F_2_s can suffer from the exposure of deleterious recessive alleles, potentially counterbalancing any heterosis. While some empirical studies support a positive correlation between heterozygosity and fitness (Springer and Stupar 2007; Yang et al. 2017), others have failed to find a consistent link (Coltman and Slate 2003; Chapman et al. 2009), suggesting that hybrid fitness may be maximized at intermediate levels of divergence, as proposed by the optimal mating distance theory (Bateson 1978; Wei & Zhang 2018). This theory posits that while genetic similarity can lead to increased homozygosity at deleterious loci, excessive divergence might disrupt co-adapted gene complexes and introduce DMIs, potentially resulting in hybrid breakdown (Barton 2001; Edmands 1999). Our findings that heterosis in survival increases with genetic distance (Fig. 3) indicates that *B. retrofracta* populations may be at an early stage of divergence, where DMIs and other genetic incompatibilities have not yet accumulated to a degree that reduces fitness. Alternatively, the disruption of coadapted positive epistatic interactions, which could inhibit fitness, may also not have diverged significantly in these populations (Lynch 1991). Nevertheless, our data suggest that parental genotype strongly influences heterosis expression, as certain parental genotypes consistently produce high-fitness offspring while others do not (Fig. 4).

We also note here that while we did not find any evidence for reproductive barriers between populations of *B. retrofracta*, multiple reproductive barriers exist between *B. retrofracta* and the more distantly related *B. stricta*, and previous research indicated that wild interspecific hybrid asexuals (including *B. stricta* x *B. retrofracta* hybrids) experienced higher rates of insect herbivory (Rushworth et al. 2020). Here, we show elevated heterozygosity in two key regions on chromosomes 1 and 7 in both wild-collected and lab-generated interspecific and intraspecific hybrids (Fig. 5), in the same approximate high-diversity regions enriched for immune-related nucleotide-binding site leucine-rich repeats (NB-LRRs) previously identified in *B. stricta* (Wang et al. 2019). A third region on chromosome 5 was similarly characterized by Wang et al. (2019). However, we found that heterozygosity was notably reduced in chromosome 5 in both wild and lab-generated *B. stricta* x *B. retrofracta* hybrids (Fig. 5).

Collectively, these findings could suggest that in *Boechera*, optimal mating distance may involve trade-offs between negative epistasis at loci related to pathogen or herbivore defense and resilience to abiotic stress.

### The influence of mating system on heterosis

While genetic divergence may explain the heterosis observed in this study, other factors, such as mating system and demographic history, must also be considered. In maize, for instance, inbreeding can purge highly deleterious alleles (Roessler et al. 2019), but weakly deleterious alleles often persist, as predicted in selfing populations (Arunkumar et al. 2015). This residual genetic load, typically referred to as inbreeding or masked load, continues to impact fitness, contributing to the frequent expression of heterosis in domesticated species (Bertorelle et al. 2022). Although heterosis is typically more pronounced in domesticated lineages (e.g. Plech et al. 2014), it also occurs in wild self-fertilizing species like *Arabidopsis*. Similarly, most *Boechera* species, including the non-hybrids in this experiment, are predominantly self-fertilizing (Roy 1995; Song et al. 2006; Rushworth 2018).

Despite the prevalence of self-fertilization, there is a striking abundance of hybrid lineages and frequent transitions to asexuality in *Boechera* (Li et al. 2017). Using genomic data, we confirmed the presence of both inter- and intraspecific hybrids, as has been shown in previous studies of *Boechera* (Schilling et al. 2018, Kantama et al. 2007). Hybrids were genomically intermediate between their parental species, with all wild-collected *B. stricta* x *B. retrofracta* hybrids appearing to be F_1_s, while wild-collected intraspecific *B. retrofracta* hybrids showed evidence of self-fertilization and/or backcrossing (Fig. 5A, B). In addition, previous studies determined that nearly all wild-collected hybrids were asexual (Beck et al. 2012, Rushworth et al. 2018, 2020). This suggests that interspecific hybrids transition to asexual reproduction concurrent with or shortly after hybridization, whereas intraspecific hybrids transition in later generations, which might be explained by different mechanisms of asexuality in *Boechera*(e.g., Windham et al. 2016: diplospory in interspecific hybrids; Carman et al. 2019: apospory in intraspecific hybrids). Determining the timing of the transition to asexuality is critical for elucidating the impact of both fitness-related factors, such as outbreeding depression or heterosis, and the nature of the alleles involved, on the formation and persistence of hybrid asexuals since they exhibit fixed traits due to a lack of recombination (Ozias-Akins & Van Dijk 2007).

### Heterosis varies with environment

Our study found that the magnitude of survival heterosis in *B. retrofracta* varied significantly between environments, with heterosis at one garden site (CAP) being three times greater than at the other (MIL). This finding is consistent with previous studies that report variability in heterosis across different environments (Bryant et al. 20017; Li et al. 2018). However, despite extensive research on the genetic basis of environmental adaptation, the specific factors influencing heterosis are still not well understood. Identifying and experimentally manipulating environmental variables could potentially enhance or reduce heterosis. For instance, during our study, poor conditions at the CAP site coincided with reduced flowering in the local plant community (C. Rushworth, pers. obs.). Biotic factors may also play a role in shaping heterosis (Wagner et al. 2021, Thompson et al. 2022), though this area remains underexplored. Future research should focus on addressing whether certain environmental variables can predict the level of heterosis, if these factors can be leveraged in controlled settings to improve fitness, and whether their effects are consistent across species or specific to particular ecosystems.

### Conclusion

Our findings offer insights into the origins and persistence of hybrids in *Boechera* populations. We propose that heterosis scales with genetic divergence in wild asexual hybrids, possibly at two key genomic regions. Masking of deleterious alleles in these regions may enhance overwinter survival. Fixation of heterosis, particularly in early generations of interspecific hybrids—where F_1_heterosis is expected to be strongest (Endler 1977)—produces highly heterotic asexual hybrids that can persist in nature. We also suggest that negative epistasis in interspecific *Boechera* hybrids contributes to reproductive isolation, with the region on chromosome 5 enriched for NB-LRRs potentially representing Dobzhansky-Muller Incompatibilities (DMIs), a phenomenon supported by studies in *Arabidopsis* (Bomblies et al. 2007). Although such DMIs may not yet have evolved among *B. retrofracta* populations, selfing could drive the accumulation of postzygotic incompatibilities (Marie-Orleach et al. 2024), and identifying the generation at which fixation occurs in asexual hybrids will clarify the nature of these DMIs (e.g., dominance and additivity). Further research is needed to uncover the genetic basis of genetic load, understand how environmental factors maintain heterosis, and explore how selective pressures influence the persistence of hybrids in natural populations.

## Supporting information

All supplemental tables

---

**Figure S1:**
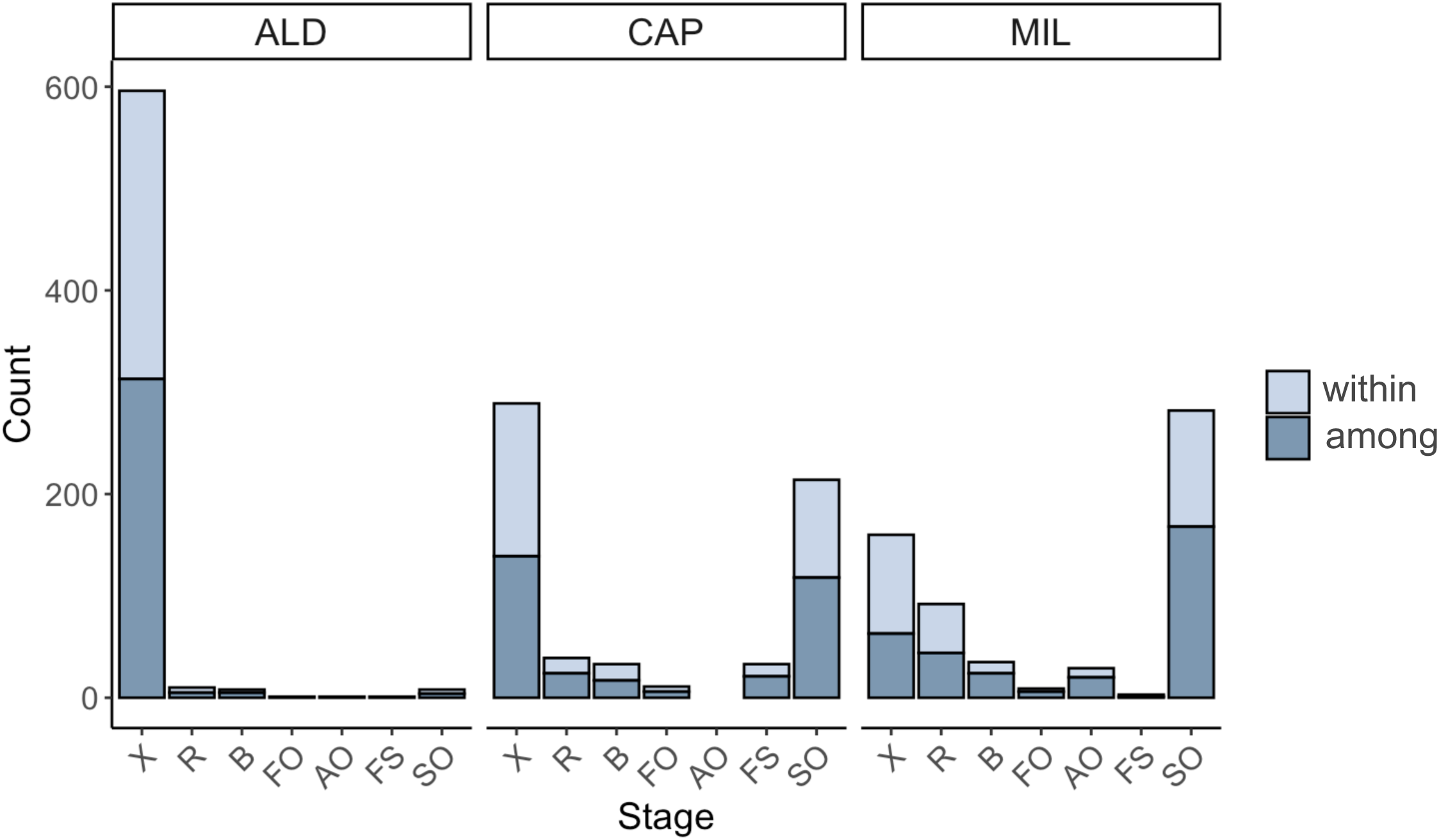
Count of offspring within (light blue) and among (dark blue) populations in three different gardens, ALD, CAP, and MIL for various life-history stages. X is died after transplanting, R is rosette, B is bolting, FO is flowers only, AO is aborted fruits only, FS is flowers and siliques, SO is siliques only. Nearly all plants in ALD garden died (X) and were removed from further analyses.

**Figure S2:**
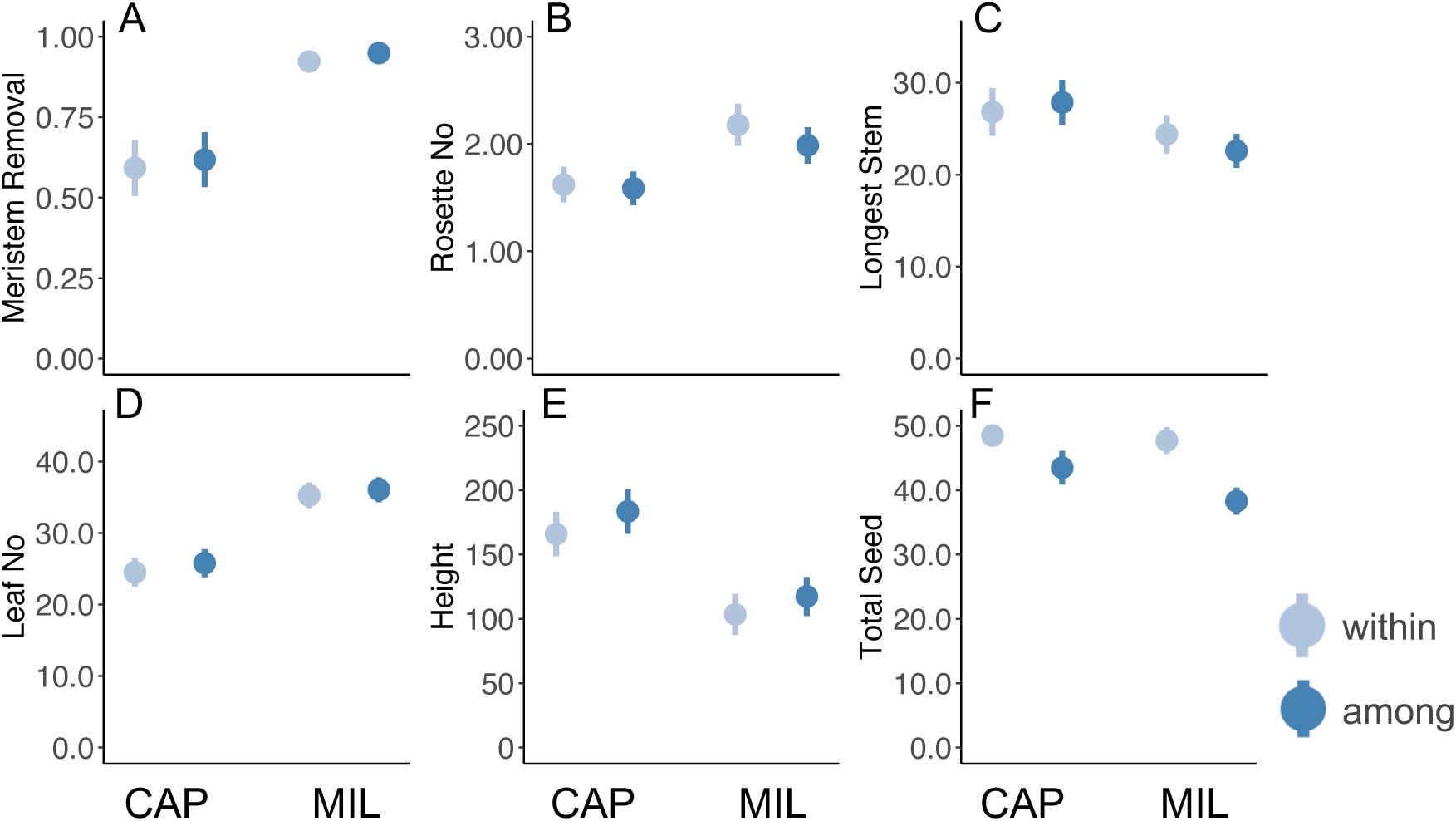
The effects of garden, cross type (within vs. among), or their interaction on fitness traits as determined by brms. Data points show conditional means of fitness for offspring from among (dark blue) and within (light blue) populations at CAP and MIL gardens. **A)** Proportion of meristem removal varies across gardens **B)** Number of rosettes per plant varies by garden. **C)** Longest stem in cm varies by garden. **D)** Number of leaves varies by garden. **E)** Plant height in cm varies by garden and cross type**. F)** Total seeds per fruit varies by the interaction between garden and cross type.

**Figure S3:**
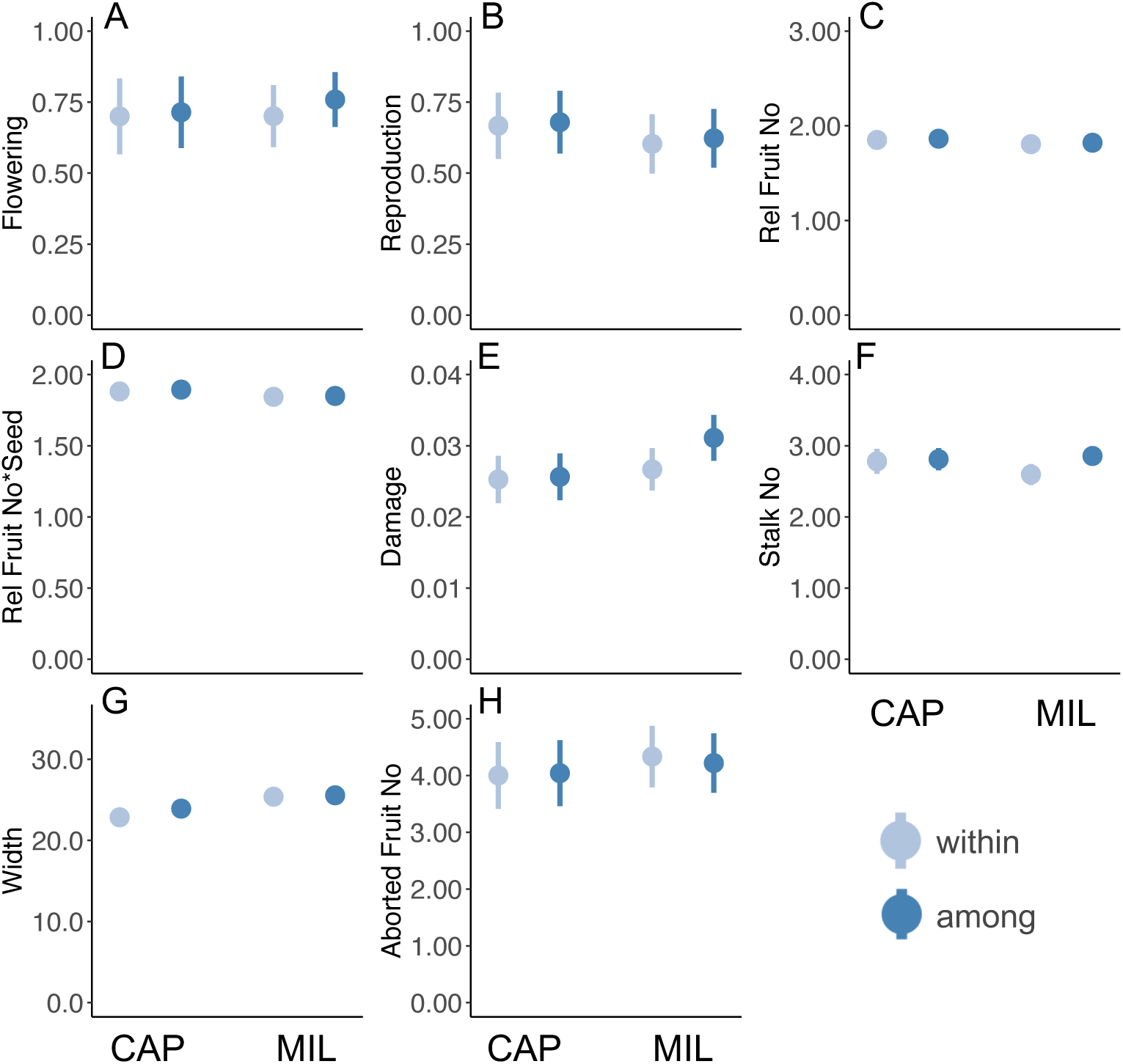
No evidence for outbreeding depression or heterosis in most fitness traits of *B. retrofracta* hybrids. Data points show conditional means of fitness for offspring from among (dark blue) and within (light blue) populations at CAP and MIL gardens. **A)** Proportion of surviving individuals that flowered. **B)** Proportion of surviving individuals that reproduced. **C)** Relative fruit number **D)** Relative fruit number * seed to calculate maximum seed set per genotype. **E)** Proportion leaf damage on total plant. **F)** Number of bolting stalks per plant was increased in among vs within only at MIL. **G)** Plant width in cm. **H)** Number of aborted fruits.

**Table S1:** Crossing combinations, including maternal and paternal sources and resulting offspring. “Among” crosses indicate crosses between individuals from different populations, while “within” crosses involve individuals from the same population. The table also includes the genetic divergence (dxy) between individuals in each cross, with a value of 0 indicating a “selfed” cross.

**Table S2:** Sample information for genomic analyses, including sample name, replicate, and species ID, mode of reproduction (sexual or asexual), number of raw sequencing reads from GBS, percent reads trimmed with CutAdapt, and corresponding SRA accession numbers.

**Table S3:** Bayesian regression models using Stan output, examining the effects of cross type (parent = par, within-population = wi, and among-population = am), garden location (CAP and MIL), and their interaction on fitness traits. The table includes standard error, 95% confidence intervals (lower and upper), potential scale reduction factor (Rhat), and bulk and tail effective sample size (ESS).

**Table S4:** Post-hoc hypothesis testing of Bayesian regression models using Stan. The table presents the effects of cross type (parent = par, within-population = wi, and among-population = am) on fitness at both garden locations (CAP and MIL). It includes standard error, 95% confidence intervals (lower and upper), and significance indication, where “NS” denotes intervals that overlap 0, and “*” indicates intervals that do not overlap 0.

**Table S5:** Bayesian regression models using Stan output, examining the effects of cross type (within-population = wi, and among-population = am -- note that here within includes selfed parents), garden location (CAP and MIL), and their interaction on fitness traits. The table includes standard error, 95% confidence intervals (lower and upper), potential scale reduction factor (Rhat), and bulk and tail effective sample size (ESS).

**Table S6:** Post-hoc hypothesis testing of Bayesian regression models using Stan. The table presents the effects of cross type (within-population = wi, and among-population = am -- note that within-population includes selfed parents) on fitness at both garden locations (CAP and MIL). The table includes standard error, 95% confidence intervals (lower and upper), and significance indication, where “NS” denotes intervals that overlap 0, and “*” indicates intervals that do not overlap 0.

**Table S7:** Post-hoc hypothesis testing of Bayesian regression models using Stan. The table presents the effects of garden (CAP and MIL) on fitness for both cross types (within-population = wi, and among-population = am -- note that within-population includes selfed parents). The table includes standard error, 95% confidence intervals (lower and upper), and significance indication, where “NS” denotes intervals that overlap 0, and “*” indicates intervals that do not overlap 0.

**Table S8:** Family means of survival estimated by Bayesian regression modeling using Stan. The table includes each unique cross identifier (cross), survival mean, standard error, and 95% confidence intervals for the CAP and MIL gardens.

**Table S9:** Parental means for each cross for survival as estimated by Bayesian regression modeling using Stan. The table includes survival means and standard errors of the maternal and paternal parent used to calculate mid-parent average.

**Table S10:** Admixture proportion (q) and interspecific ancestry (Q12) for two genetic groups within B. retrofracta and their de novo and wild-collected hybrids.

**Table S11:** Admixture proportion (q) and interspecific ancestry (Q12) for two species (B. stricta and B. retrofracta) and their hybrids (de novo and wild collected hybrids).

## References

Allard, R. W., & Bradshaw, A. D. (1964). Implications of genotype-environmental interactions in applied plant breeding.

Andrews, S. (2010). FastQC: A Quality Control Tool for High Throughput Sequence Data.

Arnold, M. L., & Martin, N. H. (2010). Hybrid fitness across time and habitats. Trends in Ecology & Evolution, 25(9), 530–536.

Arunkumar, R., Ness, R. W., Wright, S. I., & Barrett, S. C. (2015). The evolution of selfing is accompanied by reduced efficacy of selection and purging of deleterious mutations. Genetics, 199(3), 817–829.

Barton, N. H. (2001). The role of hybridization in evolution. Molecular ecology, 10(3), 551–568.

Bates, D., Maechler, M., Bolker, B., Walker, S., Christensen, R. H. B., Singmann, H.,…& Bolker, M. B. (2015). Package ‘lme4’. Convergence, 12(1), 2.

Bateson, P. (1978). Sexual imprinting and optimal outbreeding. Nature, 273(5664), 659–660.

Bateson, W. (1909). Heredity and variation in modern lights. In Darwin and Modern Science.

Beck, J. B., Alexander, P. J., Allphin, L., Al-Shehbaz, I. A., Rushworth, C., Bailey, C. D., & Windham, M. D. (2012). Does hybridization drive the transition to asexuality in diploid Boechera?. Evolution, 66(4), 985–995.

Bertorelle, G., Raffini, F., Bosse, M., Bortoluzzi, C., Iannucci, A., Trucchi, E.,…& van Oosterhout, C. (2022). Genetic load: genomic estimates and applications in non-model animals. Nature Reviews Genetics, 23(8), 492–503.

Bomblies, K., & Weigel, D. (2007). Hybrid necrosis: autoimmunity as a potential gene-flow barrier in plant species. Nature Reviews Genetics, 8(5), 382–393.

Bryant, J. R., López-Villalobos, N., Pryce, J. E., Holmes, C. W., Johnson, D. L., & Garrick, D. J. (2007). Effect of environment on the expression of breed and heterosis effects for production traits. Journal of Dairy Science, 90(3), 1548–1553.

Bürkner, P. (2017). “brms: An R Package for Bayesian Multilevel Models Using Stan.” Journal of Statistical Software, 80(1), 1–28.

Chapman, J. R., Nakagawa, S., Coltman, D. W., Slate, J., & Sheldon, B. (2009). A quantitative review of heterozygosity–fitness correlations in animal populations. Molecular ecology, 18(13), 2746–2765.

Carman, J. G., Mateo de Arias, M., Gao, L., Zhao, X., Kowallis, B. M., Sherwood, D. A.,…& Windham, M. D. (2019). Apospory and diplospory in diploid Boechera (Brassicaceae) may facilitate speciation by recombination-driven apomixis-to-sex reversals. Frontiers in Plant Science, 10, 724.

Charlesworth, B., Charlesworth, D., & Morgan, M. T. (1990). Genetic loads and estimates of mutation rates in highly inbred plant populations. Nature, 347(6291), 380–382.

Charlesworth, D., & Charlesworth, B. (1987). Inbreeding depression and its evolutionary consequences. Annual Review of Ecology and Systematics, 18, 237–268.

Charlesworth, D., Morgan, M. T., & Charlesworth, B. (1990). Inbreeding depression, genetic load, and the evolution of outcrossing rates in a multilocus system with no linkage. Evolution, 44(6), 1469–1489.

Charlesworth, D., & Willis, J. H. (2009). The genetics of inbreeding depression. Nature Reviews Genetics, 10(11), 783–796.

Cheptou, P. O., & Donohue, K. (2011). Environment-dependent inbreeding depression: its ecological and evolutionary significance. New Phytologist, 189(2), 395–407.

Corral, J. M., Vogel, H., Aliyu, O. M., Hensel, G., Thiel, T., Kumlehn, J., & Sharbel, T. F. (2013). A conserved apomixis-specific polymorphism is correlated with exclusive exonuclease expression in premeiotic ovules of apomictic Boechera species. Plant Physiology, 163(4), 1660–1672.

Coyne, J. A., & Orr, H. A. (2004). Speciation. Sunderland, MA, USA: Sinauer Associates.

Crow, J. F. (1948). Alternative hypotheses of hybrid vigor. Genetics, 33(5), 477–487.

Crow, J. F. (1952). Dominance and overdominance. In J. W. Gowen (Ed.), Heterosis (pp. 282–297). Ames: Iowa State College Press.

Crow, J. F., & Kimura, M. (1970). An Introduction to Population Genetic Theory. New York: Harper and Row.

Danecek, P., Auton, A., Abecasis, G., Albers, C. A., Banks, E., DePristo, M. A.,…& 1000 Genomes Project Analysis Group. (2011). The variant call format and VCFtools. Bioinformatics, 27(15), 2156-2158.

Darwin, C. (1877). The Effects of Cross and Self Fertilisation in the Vegetable Kingdom. New York: D. Appleton.

Dapp, M., Reinders, J., Bédiée, A., Balsera, C., Bucher, E., Theiler, G.,…& Paszkowski, J. (2015). Heterosis and inbreeding depression of epigenetic Arabidopsis hybrids. Nature Plants, 1(7), 1–8.

DeRaad, D. (2023). SNPfiltR: Interactively Filter SNP Datasets. R package version 1.0.1.

Dobzhansky, T. (1951). Genetics and the Origin of Species (3rd ed.). New York: Columbia University Press.

Duvick, J. (2001). Prospects for reducing fumonisin contamination of maize through genetic modification. Environmental health perspectives, 109(suppl 2), 337–342.

Edmands, S. (1999). Heterosis and outbreeding depression in interpopulation crosses spanning a wide range of divergence. Evolution, 53(6), 1757–1768.

Edmands, S. (2002). Does parental divergence predict reproductive compatibility? Trends in Ecology & Evolution, 17(11), 520–527.

Endler, J. A. (1977). Geographic variation, speciation, and clines (No. 10). Princeton University Press.

Falconer, D. S. (1996). Introduction to Quantitative Genetics (4th ed.). Essex: Longman.

Glémin, S. (2003). How are deleterious mutations purged? Drift versus nonrandom mating. Evolution, 57(12), 2678–2687.

Haldane, J. B. S. (1937). The effect of variation of fitness. The American Naturalist, 71(735), 337–349.

Hamilton, M. B., & Mitchell-Olds, T. (1994). The mating system and relative performance of selfed and outcrossed progeny in Arabis fecunda (Brassicaceae). American Journal of Botany, 81(10), 1252–1256.

Huang, X., Yang, S., Gong, J., Zhao, Q., Feng, Q., Zhan, Q.,…& Han, B. (2016). Genomic architecture of heterosis for yield traits in rice. Nature, 537(7622), 629–633.

Husband, B. C., & Schemske, D. W. (1996). Evolution of the magnitude and timing of inbreeding depression in plants. Evolution, 50(1), 54–70.

Kantama, L., Sharbel, T. F., Schranz, M. E., Mitchell-Olds, T., de Vries, S., & de Jong, H. (2007). Diploid apomicts of the Boechera holboellii complex display large-scale chromosome substitutions and aberrant chromosomes. Proceedings of the National Academy of Sciences, 104(35), 14026–14031.

Kimura, M., Maruyama, T., & Crow, J. F. (1963). The mutation load in small populations. Genetics, 48(10), 1303.

Korunes, K. L., & Samuk, K. (2021). pixy: Unbiased estimation of nucleotide diversity and divergence in the presence of missing data. Molecular Ecology Resources, 21(4), 1359–1368.

Krieger, U., Lippman, Z. B., & Zamir, D. (2010). The flowering gene SINGLE FLOWER TRUSS drives heterosis for yield in tomato. Nature genetics, 42(5), 459–463.

Lande, R., & Schemske, D. W. (1985). The evolution of self-fertilization and inbreeding depression in plants. I. Genetic models. Evolution, 39(1), 24–40.

Lee, C. R., & Mitchell-Olds, T. (2011). Quantifying effects of environmental and geographical factors on patterns of genetic differentiation. Molecular Ecology, 20(22), 4631–4642.

Li, H. (2013). Aligning sequence reads, clone sequences and assembly contigs with BWA-MEM. arXiv preprint arXiv:1303.3997.

Li, H., Handsaker, B., Wysoker, A., Fennell, T., Ruan, J., Homer, N.,…& 1000 Genome Project Data Processing Subgroup. (2009). The sequence alignment/map format and SAMtools. Bioinformatics, 25(16), 2078-2079.

Li, H., & Durbin, R. (2009). Fast and accurate short read alignment with Burrows–Wheeler transform. Bioinformatics, 25(14), 1754–1760.

Li, F. W., Rushworth, C. A., Beck, J. B., & Windham, M. D. (2017). Boechera microsatellite website: an online portal for species identification and determination of hybrid parentage. Database, 2017, baw169.

Li, Z., Coffey, L., Garfin, J., Miller, N. D., White, M. R., Spalding, E. P.,…& Hirsch, C. N. (2018). Genotype-by-environment interactions affecting heterosis in maize. PloS one, 13(1), e0191321.

Li, Z., Sun, J., & Hirsch, C. N. (2022). Understanding environmental modulation of heterosis. Plant Breeding Reviews, 46, 219–237.

Liberatore, K. L., Jiang, K., Zamir, D., & Lippman, Z. B. (2013). Heterosis: The case for single-gene overdominance. Polyploid and hybrid genomics, 137-152.

Lynch, M. (1991). The genetic interpretation of inbreeding depression and outbreeding depression. Evolution, 45(3), 622–629.

Lynch, M., Conery, J., & Bürger, R. (1995). Mutational meltdowns in sexual populations. Evolution, 49(6), 1067–1080.

Lynch, M., & Gabriel, W. (1990). Mutation load and the survival of small populations. Evolution, 44(7), 1725–1737.

Lynch, M., & Walsh, B. (1998). Genetics and Analysis of Quantitative Traits.

Marie-Orleach, L., Glémin, S., Brandrud, M. K., Brysting, A. K., Gizaw, A., Gustafsson, A. L. S.,…& Birkeland, S. (2024). How Does Selfing Affect the Pace and Process of Speciation?. Cold Spring Harbor Perspectives in Biology, a041426.

Martin, M. (2011). Cutadapt removes adapter sequences from high-throughput sequencing reads. EMBnet.journal, 17(1), 10–12.

Mau, M., Corral, J. M., Vogel, H., Melzer, M., Fuchs, J., Kuhlmann, M.,…& Sharbel, T. F. (2013). The conserved chimeric transcript UPGRADE2 is associated with unreduced pollen formation and is exclusively found in apomictic Boechera species. Plant Physiology, 163(4), 1640–1659.

Mau, M., Liiving, T., Fomenko, L., Goertzen, R., Paczesniak, D., Böttner, L., & Sharbel, T. F. (2021). The spread of infectious asexuality through haploid pollen. New Phytologist, 230(2), 804–820.

McKenna, A., Hanna, M., Banks, E., Sivachenko, A., Cibulskis, K., Kernytsky, A.,…& DePristo, M. A. (2010). The Genome Analysis Toolkit: a MapReduce framework for analyzing next-generation DNA sequencing data. Genome Research, 20(9), 1297–1303.

Mercer, K. L., Andow, D. A., Wyse, D. L., & Shaw, R. G. (2007). Stress and domestication traits increase the relative fitness of crop–wild hybrids in sunflower. Ecology Letters, 10(5), 383–393.

Meyer, R. C., Kusterer, B., Lisec, J., Steinfath, M., Becher, M., Scharr, H.,…& Altmann, T. (2010). QTL analysis of early stage heterosis for biomass in Arabidopsis. Theoretical and Applied Genetics, 120, 227–237.

Muller, H. J. (1942). Isolating mechanisms, evolution, and temperature. In Biological Symposia (Vol. 6, pp. 71-125). Lancaster, PA: Jaques Cattell Press.

Oakley, C. G., & Winn, A. A. (2012). Effects of population size and isolation on heterosis, mean fitness, and inbreeding depression in a perennial plant. New Phytologist, 196(1), 261–270.

Oakley, C. G., Lundemo, S., Ågren, J., & Schemske, D. W. (2019). Heterosis is common and inbreeding depression absent in natural populations of Arabidopsis thaliana. Journal of evolutionary biology, 32(6), 592–603.

Ohta, T. (1973). Slightly deleterious mutant substitutions in evolution. Nature, 246(5428), 96–98.

Ozias-Akins, P., & van Dijk, P. J. (2007). Mendelian genetics of apomixis in plants. Annu. Rev. Genet., 41(1), 509–537.

Paina, C., Byrne, S. L., Studer, B., Rognli, O. A., & Asp, T. (2016). Using a candidate gene-based genetic linkage map to identify QTL for winter survival in perennial ryegrass. PLoS One, 11(3), e0152004.

Plech, M., de Visser, J. A. G., & Korona, R. (2014). Heterosis is prevalent among domesticated but not wild strains of Saccharomyces cerevisiae. G3: Genes, Genomes, Genetics, 4(2), 315-323.

Prasad, K. V., Song, B. H., Olson-Manning, C., Anderson, J. T., Lee, C. R., Schranz, M. E.,…& Mitchell-Olds, T. (2012). A gain-of-function polymorphism controlling complex traits and fitness in nature. science, 337(6098), 1081–1084.

R Core Team (2021). R: A language and environment for statistical computing. R Foundation for Statistical Computing, Vienna, Austria. URL https://www.R-project.org/.

Ralls, K., Ballou, J. D., & Templeton, A. (1988). Estimates of lethal equivalents and the cost of inbreeding in mammals. Conservation Biology, 2(2), 185–193.

Roessler, K., Muyle, A., Diez, C. M., Gaut, G. R., Bousios, A., Stitzer, M. C.,…& Gaut, B. S. (2019). The genome-wide dynamics of purging during selfing in maize. Nature plants, 5(9), 980–990.

Roy, B. A. (1995). The breeding systems of six species of Arabis (Brassicaceae). American Journal of Botany, 82(7), 869–877.

Rushworth, C. A., Windham, M. D., Keith, R. A., & Mitchell-Olds, T. (2018). Ecological differentiation facilitates fine-scale coexistence of sexual and asexual *Boechera*. American Journal of Botany, 105(12), 2051–2064.

Rushworth, C. A., & Mitchell-Olds, T. (2021). The evolution of sex is tempered by costly hybridization in *Boechera*(rock cress). Journal of Heredity, 112(1), 67–77.

Rushworth, C. A., Brandvain, Y., & Mitchell-Olds, T. (2020). Identifying the fitness consequences of sex in complex natural environments. Evolution Letters, 4(6), 516–529.

Rushworth, C. A., Song, B. H., Lee, C. R., & Mitchell-Olds, T. (2011). *Boechera*, a model system for ecological genomics. Molecular Ecology, 20(23), 4843–4857.

Rushworth, C. A., Wagner, M. R., Mitchell-Olds, T., & Anderson, J. T. (2022). The *Boechera* model system for evolutionary ecology. American Journal of Botany, 109(11), 1939–1961.

Schemske, D.W., 1983. Breeding system and habitat effects on fitness components in three neotropical Costus (Zingiberaceae). Evolution, pp.523–539.

Schilling, M. P., Gompert, Z., Li, F. W., Windham, M. D., & Wolf, P. G. (2018). Admixture, evolution, and variation in reproductive isolation in the Boechera puberula clade. BMC evolutionary biology, 18, 1–14.

Schoen, D. J. (1983). Relative fitnesses of selfed and outcrossed progeny in Gilia achilleifolia (Polemoniaceae). Evolution, 292-301.

Schranz, M. E., Dobeš, C., Koch, M. A., & Mitchell-Olds, T. (2005). Sexual reproduction, hybridization, apomixis, and polyploidization in the genus *Boechera* (Brassicaceae). American Journal of Botany, 92(11), 1797–1810.

Schranz, M. E., Kantama, L., De Jong, H., & Mitchell-Olds, T. (2006). Asexual reproduction in a close relative of *Arabidopsis*: a genetic investigation of apomixis in *Boechera* (Brassicaceae). New Phytologist, 171(2), 425–438.

Shastry, V., Adams, P. E., Lindtke, D., Mandeville, E. G., Parchman, T. L., Gompert, Z., & Buerkle, C. A. (2021). Model-based genotype and ancestry estimation for potential hybrids with mixed-ploidy. Molecular Ecology Resources, 21(5), 1434–1451.

Shull, G. H. (1952). Beginnings of the heterosis concept. In J. W. Gowen (Ed.), Heterosis (pp. 31–33). Ames: Iowa State College Press.

Sobel, J. M., Chen, G. F., Watt, L. R., & Schemske, D. W. (2010). The biology of speciation. Evolution, 64(2), 295–315.

Song, B. H., Clauss, M. J., Pepper, A., & Mitchell-Olds, T (2006). Geographic patterns of microsatellite variation in *Boechera stricta*, a close relative of *Arabidopsis*. Molecular Ecology, 15(2), 357–369.

Song, B. H., & Mitchell-Olds, T (2007). High genetic diversity and population differentiation in *Boechera fecunda*, a rare relative of *Arabidopsis*. Molecular Ecology, 16(19), 4079–4088.

Soto, T. Y., Rojas-Gutierrez, J. D., & Oakley, C. G. (2023). Can heterosis and inbreeding depression explain the maintenance of outcrossing in a cleistogamous perennial?. American Journal of Botany, 110(10), e16240.

Springer, N. M., & Stupar, R. M. (2007). Allelic variation and heterosis in maize: how do two halves make more than a whole?.Genome research, 17(3), 264–275.

Stelkens, R. B., Schmid, C., & Seehausen, O. (2015). Hybrid breakdown in cichlid fish. PloS one, 10(5), e0127207.

Sweigart, A. L., Fishman, L., & Willis, J. H. (2006). A simple genetic incompatibility causes hybrid male sterility in Mimulus. Genetics, 172(4), 2465–2479.

Thompson, K. A., Brandvain, Y., Coughlan, J. M., Delmore, K. E., Justen, H., Linnen, C. R.,…& Stelkens, R. (2023). The ecology of hybrid incompatibilities. Cold Spring Harbor Perspectives in Biology, a041440.

Thompson, K. A., Peichel, C. L., Rennison, D. J., McGee, M. D., Albert, A. Y., Vines, T. H.,…& Schluter, D. (2022). Analysis of ancestry heterozygosity suggests that hybrid incompatibilities in threespine stickleback are environment dependent. PLoS biology, 20(1), e3001469.

Wagner, M. R., Tang, C., Salvato, F., Clouse, K. M., Bartlett, A., Vintila, S.,…& Kleiner, M. (2021). Microbe-dependent heterosis in maize. Proceedings of the National Academy of Sciences, 118(30), e2021965118.

Wang, B., Mojica, J. P., Perera, N., Lee, C. R., Lovell, J. T., Sharma, A.,…& Mitchell-Olds, T. (2019). Ancient polymorphisms contribute to genome-wide variation by long-term balancing selection and divergent sorting in Boechera stricta. Genome biology, 20, 1–15.

Wang, H., Su, B., Zhang, Y., Shang, M., Li, S., Xing, D.,…& Wang, X. (2024). From heterosis to outbreeding depression: genotype-by-environment interaction shifts hybrid fitness in opposite directions. Genetics, iyae090.

Wei, X., & Zhang, J. (2018). The optimal mating distance resulting from heterosis and genetic incompatibility. Science Advances, 4(11), eaau5518.

Whitlock, M. C., Ingvarsson, P. K., & Hatfield, T. (2000). Local drift load and the heterosis of interconnected populations. Heredity, 84(4), 452–457.

Wickham, H. (2011). ggplot2. Wiley Interdisciplinary Reviews: Computational Statistics, 3(2), 180–185.

Willi, Y., & Van Buskirk, J. (2005). Genomic compatibility occurs over a wide range of parental genetic similarity in an outcrossing plant. Proceedings of the Royal Society B: Biological Sciences, 272(1570), 1333–1338.

Windham, M. D., & Al-Shehbaz, I. A. (2006). New and noteworthy species of *Boechera* (Brassicaceae) I: sexual diploids. Harvard Papers in Botany, 11(1), 61–88.

Windham, M. D., Beck, J. B., Li, F. W., Allphin, L., Carman, J. G., Sherwood, D. A.,…& Al-Shehbaz, I. A. (2016). Searching for diamonds in the apomictic rough: a case study involving oechera lignifera (Brassicaceae). Systematic Botany, 40(4), 1031–1044.

Yang, M., Wang, X., Ren, D., Huang, H., Xu, M., He, G., & Deng, X. W. (2017). Genomic architecture of biomass heterosis in Arabidopsis. Proceedings of the National Academy of Sciences, 114(30), 8101–8106.

Zuellig, M. P., & Sweigart, A. L. (2018). Gene duplicates cause hybrid lethality between sympatric species of Mimulus. PLoS genetics, 14(4), e1007130.

